# Synthetic RNA-protein decoy granules to prevent SARS-CoV-2 infection

**DOI:** 10.64898/2026.01.16.699918

**Authors:** Or Willinger, Naor Granik, Tai Salomon, Sarah Goldberg, Roee Amit

## Abstract

Synthetic receptor decoys offer a promising strategy to block viral entry but are often limited by instability and rapid clearance. Here we introduce AntiCoV decoy particles: phase-separated synthetic RNA-protein nanoparticles that display multivalent human ACE2 and act as robust decoy receptors for SARS-CoV-2. The granules remain structurally stable for at least two weeks at room temperature. Using confocal imaging and FRET, we show that the SARS-CoV-2 receptor-binding domain is absorbed into the granule matrix with nanometer-scale proximity to ACE2. In viral entry assays with Delta and Omicron (BA.1) variants, AntiCoV decoy particles achieve complete inhibition of infection at low micromolar concentrations and outperform soluble ACE2, which also exhibits infection enhancement at low doses. A minimal kinetic model suggests a multistep spike-priming mechanism that reconciles this biphasic response. AntiCoV decoy particles therefore provide a stable, modular, and pan-variant antiviral platform with potential for low-cost, room-temperature-stable prophylaxis.

## Introduction

The recent corona virus pandemic has brought to light the need to expand our repertoire of antiviral treatment. Antiviral therapeutics generally fall into two broad categories. The first includes replication inhibitors, small molecules that target viral enzymes or structural proteins essential for the replication cycle[1]. Examples include protease inhibitors (e.g., Paxlovid[2]), replicase inhibitors (e.g., Remdesivir[3], Molnupiravir[4], Acyclovir[5]), and agents that target envelope proteins (e.g., Umifenovir[6], Tamiflu[7]). These drugs are typically virus-specific, and efforts to repurpose them across viral families have met with limited success, although high-throughput screens of drug libraries keep improving[8,9]. While nucleotide analogs offer broad-spectrum potential, their similarity to host nucleotides makes them prone to off-target effects, leading to cytotoxicity via disruption of host polymerases activity[5].

The second category comprises immunogenic agents, such as vaccines and monoclonal antibody therapies. To date, vaccines remain the most effective prophylactic intervention against infectious diseases and have eradicated or contained deadly pathogens such as smallpox[10], measles[11], and polio[12]. However, logistical limitations – such as the need for ultra-cold storage of mRNA vaccines[13] – pose significant challenges to equitable distribution, especially in low-resource settings, as demonstrated during the COVID-19 pandemic[14,15]. Monoclonal antibodies provide post-exposure treatment by targeting viral proteins with high specificity, yet their efficacy diminishes in the face of viral mutations, with a single amino acid change sometimes sufficient to confer resistance, as seen with SARS-CoV-2 variants and the rapid obsolescence of antibody cocktails (e.g. Regeneron)[16]. In general, high specificity often translates into high susceptibility to viral evolution.

One underexplored frontier in antiviral defense is the utilization of decoy strategies for prophylactic protection. Soluble isoforms or recombinant versions of natural human proteins that viruses employ as receptors have been demonstrated in the past to inhibit viral infections[17–22]. However, untethered from their membranal component, these decoy receptors – natural or recombinant – tend to have a short half-life[23], and therefore are often fused to stabilizing domains, more commonly the Fc region of IgG antibodies[18,21,24,25]. Fc regions were shown to sometimes elicit antibody-dependent enhancement[19], where a virus which was bound by an Fc-fused receptor is endocytosed into immune cells, and subsequently triggers inflammatory pathways and cell death[19]. As a result, despite their potential to be both a highly specific antiviral therapeutic and one which is insensitive to viral mutations, soluble receptor-based antiviral therapeutics have yet to reach the clinic.

We previously developed synthetic long non-coding RNA (slncRNA) constructs capable of undergoing phase separation to form stable gel-like condensates called sRNP granules when bound by the cognate RNA-binding tandem dimer of the PP7 phage coat protein (tdPCP)[26,27]. These particles can be functionalized by fusing recombinant protein cargos to the tdPCP component, enabling modular targeting[26–28]. In a proof-of-concept study, granules loaded with recombinant human ACE2 sequestered SARS-CoV-2 spike-mimicking beads, effectively acting as decoy “sponges” for virions [29]. Here, we expand on these findings to demonstrate that granules infused with the hACE2 extracellular domain are stable over a long period of time at room temperature, and also generate robust antiviral activity in entry assays for the two SARS2 variants Delta and Omicron (BA.1). Our findings pose a potential solution to the stability problem of decoy proteins and may thus provide a pathway forward to finally bringing these important potential therapeutic proteins to the clinic (Figure 1A).

**Figure 1:**
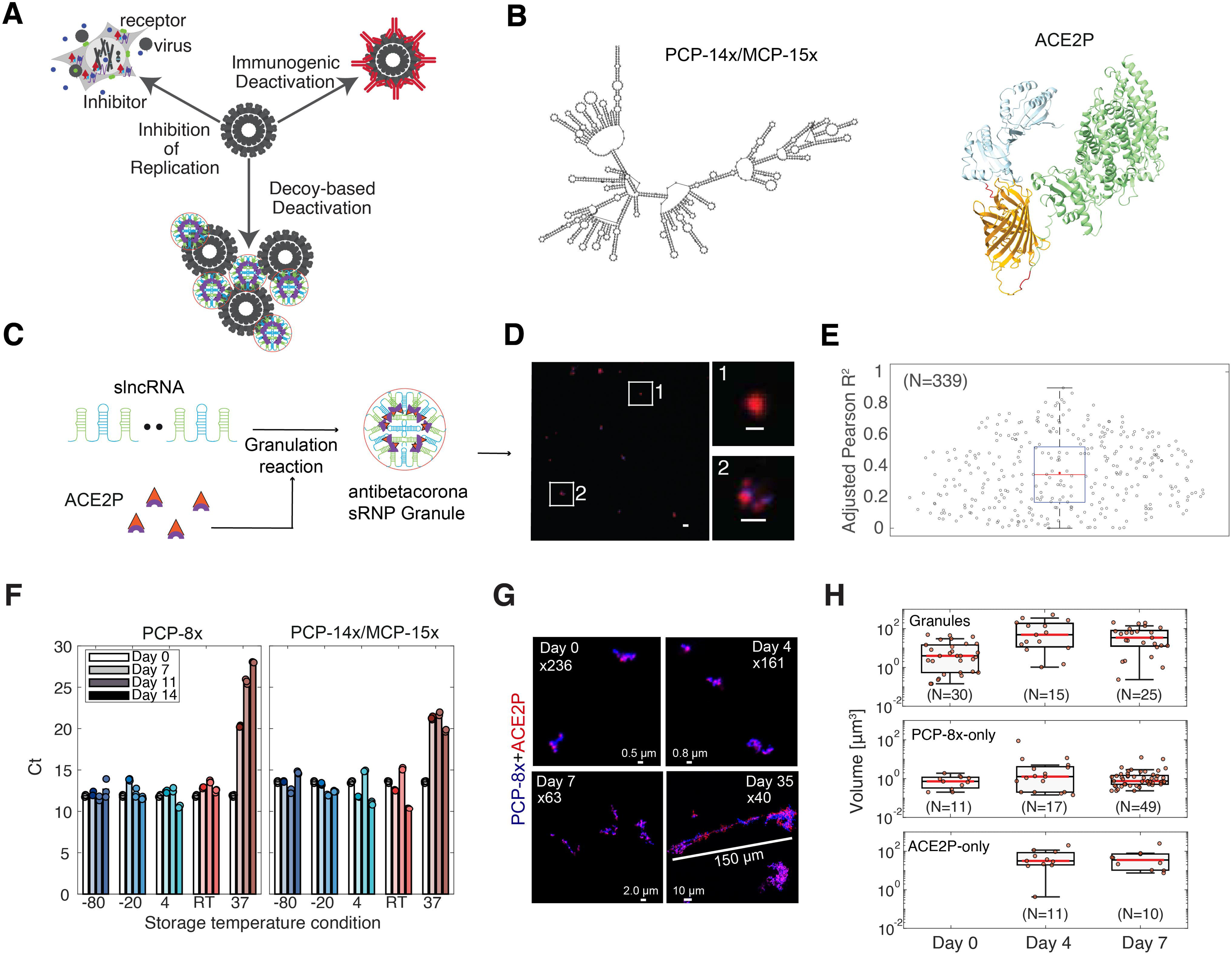
Anti-SARS-CoV-2 SRNP granules as viral decoys. **(A)** Schematic showing the three antiviral therapeutic approaches. (Upper-left) replication prevention. (Upper-right) immunogenic marking via antibodies for deactivation by the immune system. (Bottom) viral inactivation via the decoy mechanism. **(B)** (Left) Model structure computed using Vienna of a 1269 nt slncRNA encoding 14 and 15 non-sequence repeating PCP and MCP hairpins, respectively. (Right) tdPCP (blue) fused to an mCherry fluorescent protein (orange) fused to ACE2 (aa1-740) encoding both a red fluorescent and anti-SARS-CoV-2 functionality. Structure was generated using AlphaFold. **(C)** (Left) sRNP granule structure formed by the binding of the hairpin binding protein (tdPCP) to the hairpins and hairpin-hairpin cross-linking. The resultant structure (middle-schema) is a gel-like nanoparticle with a semipermeable slncRNA shell encasing a high density of ∼100 hairpin-binding functional proteins[26]. **(D)** (Left image) Representative image of ACE2 granules in a typical field of view. Scalebar: 5µm. (Right images) Enlarged images highlighted by white squares on the left. (Right-top) protein only condensate. (Right-bottom) An sRNP granule of protein co-localized with slncRNA. Scalebar of enlarged images 1µm and 2µm for top and bottom, respectively. (E) Pearson correlation between mCherry and AF405 signals intensities of 339mCherry-positive events. Representative events – see 1 and 2 from panel B and Figure S2A. (F) qPCR mean Ct values of slncRNA-only granules stored in various temperatures over the course of a week. **(G)** Representative events depicting aging and growth of ACE2 decoy particles over the course of 35 days. Red – ACE2P. Blue – PCP 8x slncRNA. **(H)** Volume calculations of AF405- and/or mCherry-positive events. A student’s t-test was used for statistical significance between each two boxplot distributions. Statistical significance was only observed for the Granules panel (top) with p <0.05 and p<0.001 for the difference between day 4 to day 0 and day 7 to day 0, respectively.

## Results

### AntiCoV granules are stable at room temperature for at least a week

We hypothesized that sRNP granules can be adapted to form stable antiviral decoy nanoparticles by fusing the tdPCP moiety to the soluble receptor domain of a virus target receptor, and that such particles can be screened for antiviral functionality via changes to the phase-separated granule structure when exposed to the viral receptor binding moiety (Figure 1A). To test our strategy for developing antiviral granules, we used our putative AntiCoV granules (reported previously[29]) composed of the ectodomain of hACE2-mCherry-tdPCP (ACE2P – Figure 1B-right) fusion protein and a synthetic long non-coding RNA molecules encoding either 8 PCP binding sites (slncRNA-PCP-8x, Figure S1A) or 14 PCP binding sites (slncRNA-PCP-14x/MCP-15x, Figure 1B-left).

We first opted to characterize the granule’s structure and long-term stability. To do so we employed a dual-label fluorescence microscopy approach with the slncRNA labelled with AF405-Uracil during synthesis, together with the ACE2-mCherry-tdPCP moiety (Figure 1C-D and Figure S1B). The image shows formation of either protein condensates (red) with faint slncRNA signal, or co-localized protein and slncRNA spots (blue), indicating formation of protein-RNA biocondensates or granules. We then computed the Pearson correlation for the red and blue intensities of 339 co-localized structures over 27 field-of-views (FoVs) and found various correlation values between red and blue intensities (Figure 1E), providing support for the formation of granulated structure with a range of protein-to-slncRNA ratios.

Next, to verify granule integrity for antiviral testing, we characterized them for structural stability and aging. To do so, we first incubated them at −80°C, −20°C, 4°C, room temperature (RT), and 37°C for two weeks and performed qPCR measurements on the slncRNA cassettes 1h post-formation (Day 0) and on days 7, 11, and 14 post-formation (Figure 1F and Figure S1C). The results show that up to RT, slncRNA levels remain relatively constant in the tested temperatures. At 37°C incubation conditions, both cassettes seem to degrade at an exponential rate for the first week. For the second week, however, the PCP-8x cassette continues this trend, while the slncRNA levels measured for PCP-14x/MCP-15x granules seem to reach a steady-state level. This result is consistent with our previous report that hetero-hairpin slncRNA cassettes showed increased cross-linking and denser packing in its granule phase as compared with homo-hairpin granules[26]. Importantly, attempts to reverse-transcribe either of the slncRNA cassettes were unsuccessful without first diluting the granules 10-fold (see Methods), and even 5min incubation at 95°C did not permit cDNA-synthesis without dilution. Polyethylene glycol (PEG) is reportedly beneficial for cDNA synthesis, even at a 12% concentration[33,34], and yet our results suggest that no cDNA was produced in a granule buffer with 10% PEG. We have previously reported that our granules are resistant to RNase A treatment[26], and the inability of a reverse transcriptase to use these cassettes as substrates implies that without a cognate RNA-binding domain, proteins are unable to interact with or infiltrate our granules.

To provide further support for the stability observed in the qPCR measurements, we imaged labelled PCP-8x-ACE2P granules that were formed and maintained at room temperature on similar time points as the qPCR measurements (Figure 1G-H and Figure S1D), and in addition also after 35 days. The results (Figure 1F) show that ACE2P-granules display co-localized clusters of protein (red) and slncRNA (blue) for all time points. The size of the structures seems to increase from ∼0.5 μm initially to ∼1 μm objects by day 4, ∼2 μm structures or larger by day 7, and an extended microcluster of ∼100 μm by day 35. This result indicates that ACE2 granules seem to coalesce over time in a process reminiscent of Ostwald ripening, maintaining what appears to be an increasing amount of RNA and a large concentration of active fluorescent protein even after 35 days. By contrast, the slncRNA-only sample (Figure S1D) does not exhibit this behavior. This result is further supported by direct granule volume measurements (Figure 1H), where only ACE2P granules exhibit volumetric increase after 4 and 7 days (Figure 1H-top), while slncRNA-only (Figure 1H-middle) and protein-only samples (Figure 1H-bottom) do not. Consequently, the ACE2P granules have the capability to store the potential decoy proteins for several weeks in room temperature consistent with therapeutic requirements for formulation stability.

### Phase-separation confirms functionality of AntiCoV decoy granules

To test for granule functionality, we incubated the putative AntiCoV decoy particles with the SARS-CoV-2 RBD fused to super-folder GFP (sfGFP) and imaged the resultant granule structure using three separate color channels (RGB) on Zeiss LSM710 confocal microscope (Figure 2A-B and Figure S2A-left). A close examination of the structures shows (Figure 2B see structures 1 and 2 to the right) dominant bright green patches with very little red and blue intermixed. In particular, the small red patches did not co-localize with the green patches but rather were found to be immediately adjacent. Due to the observed discontinued strong brightness in the green patches, we hypothesized that we may be observing patches of Forster resonance energy transfer (FRET) between the RBD-sfGFP and ACE2-mCherry-tdPCP. To check for green-red FRET, we rearranged our filters to capture red channel emissions while exciting the sfGFP (a built-in mode within the LSM710 confocal microscope). The resultant images (Figure 2C) exhibit a strong FRET intensity signal all across the structures, indicating strong binding and close proximity of the RBD to the ACE2. We then analyzed the FRET efficiency across our structures and found a value of 0.2 to 0.3, corresponding to an average molecular separation of 6-8 nm between the fluorescent moieties of the RBD and ACE2 molecules (Figure 2D). We next studied the FRET signal that resulted from gramules incubated with an increasing concentration of RBD-sfGFP (Figure 2E). We found a small amount of FRET signal for low concentrations (below 0.0685 mg_RBD_/ml). However, a strong FRET signal emerged (i.e. above sfGFP-only basal FRET intensity) when the RBD:ACE2 molecular ratio was 3:2 (0.0685 mg_RBD_/ml). Interestingly, this trimer:dimer ratio is the natural state of binding of SARS-CoV-2 spike protein and ACE2[26]. Since in this case the RBD-sfGFP does not contain the heptad repeat domains required for spike trimerization, our results suggest that optimal spike-ACE2 interaction, while strongest at a ratio of three RBDs to two ACE2, is not necessarily related to spike trimerization.

**Figure 2-.**
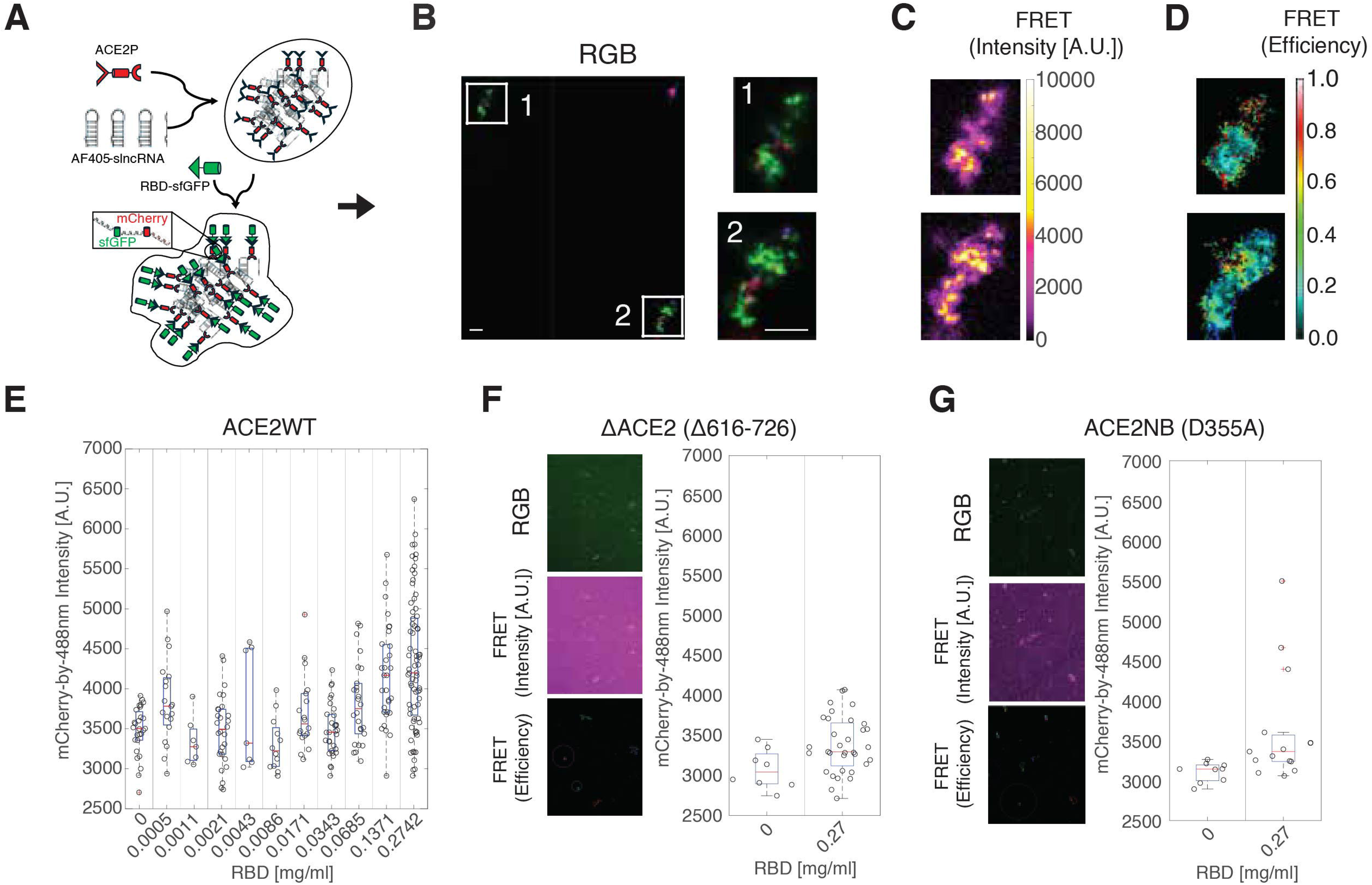
ACE2 decoy granules exhibit FRET excitation with SARS-CoV-2 RBD. **(A)** Scheme of assay: ACE2 labelled with an mCherry fluorescent protein and fused to tdPCP is incubated with a slncRNA cassette, encoding for multiple PCP-binding hairpins. 1h post incubation, an sfGFP-labelled RBD is added, facilitating FRET with the mCherry-labelled ACE2 proteins bound to the slncRNA granules. **(B)** (Left) Representative image of ACE granules together with RBD (green label). Scalebar: 5µm. (Right) Enlarged images highlighted by white squares on the left showing typical triple labelled structures. Scalebar: 5µm. **(C-D)** FRET intensity signal **(C)** and FRET efficiency **(D)** measured for the highlighted granule structures shown in Figure 2B. (**E)** Putative green-red FRET excitation events obtained for various RBD concentrations. **(F-G)** (Left) Images of triple-labelled granules (top), FRET intensity (middle), and FRET efficiency (bottom), and (right) putative intensity of FRET excitation events obtained for (**F**) the non-dimerizing ACE2 (Δ616-726) and (**G**) the non-RBD-binding missense ACE2 mutant (D355A).

To provide additional support for our findings regarding RBD-ACE2 interaction and to test for specificity, we prepared two mutant ACE2 moieties: a variant (ΔACE2) that lacks the “neck” domain responsible for dimerization[31] (Figure 2F and Figure S2A-middle), and a variant with a missense mutation (ACE2NB) that abolishes RBD binding[30] (Figure 2G and Figure S2A-right). The two mutant ACE2 variants formed tightly bound granules (Figure S2B), as was observed for the dimerized ACE2, but we did not observe colocalization of RBD-sfGFP, nor any meaningful FRET signal. Consequently, we observe a strong ACE2-spike interaction within the the granules that is dependent on dimerization of ACE2 and the binding of at least three RBD moieties.

### ACE2 decoy particles fully inhibit SARS-CoV-2 infection and suggests a more complex infection mechanism

We next wanted to test the efficacy of ACE2 decoy granules against both the Delta and Omicron BA.1 SARS-CoV-2 variants via a viral entry assay using Vero E6 cells (see Figure 3A and Methods). An initial scan of the infected cells revealed that AntiCoV decoy granules demonstrated complete inhibition of infection at the highest granule concentrations (Figure 3B). A closer observation of the anti-viral response (Figure 3C-D) further revealed enhancement of infection at lower protein concentration, which transitioned to complete inhibition of infection at ACE2 concentrations >3μg/ml. In particular, the protein-only samples showed stronger enhancement (empty circles) compared with the decoy granules (filled circles) and transitioned to inhibition at higher protein concentration thresholds. This difference can be quantified by computing the IC_50_ value for both the granules and protein-only cases (Figure S3). For the Delta variant, a 7-fold reduction (∼87%) in IC_50_ was recorded (3.15µg/ml and 22.9µg/ml, for the decoy granules and ACE2-only, respectively) while a ∼33% reduction was recorded for the Omicron BA.1 variant (IC_50_ = 3.1µg/ml and 4.6µg/ml for decoy granules and ACE2-only, respectively). Importantly, a slncRNA-only control elicited no antiviral response at any concentration for either variant, and a weak, decoy-like response at the highest concentrations (Figure 3C-D, red).

**Figure 3-.**
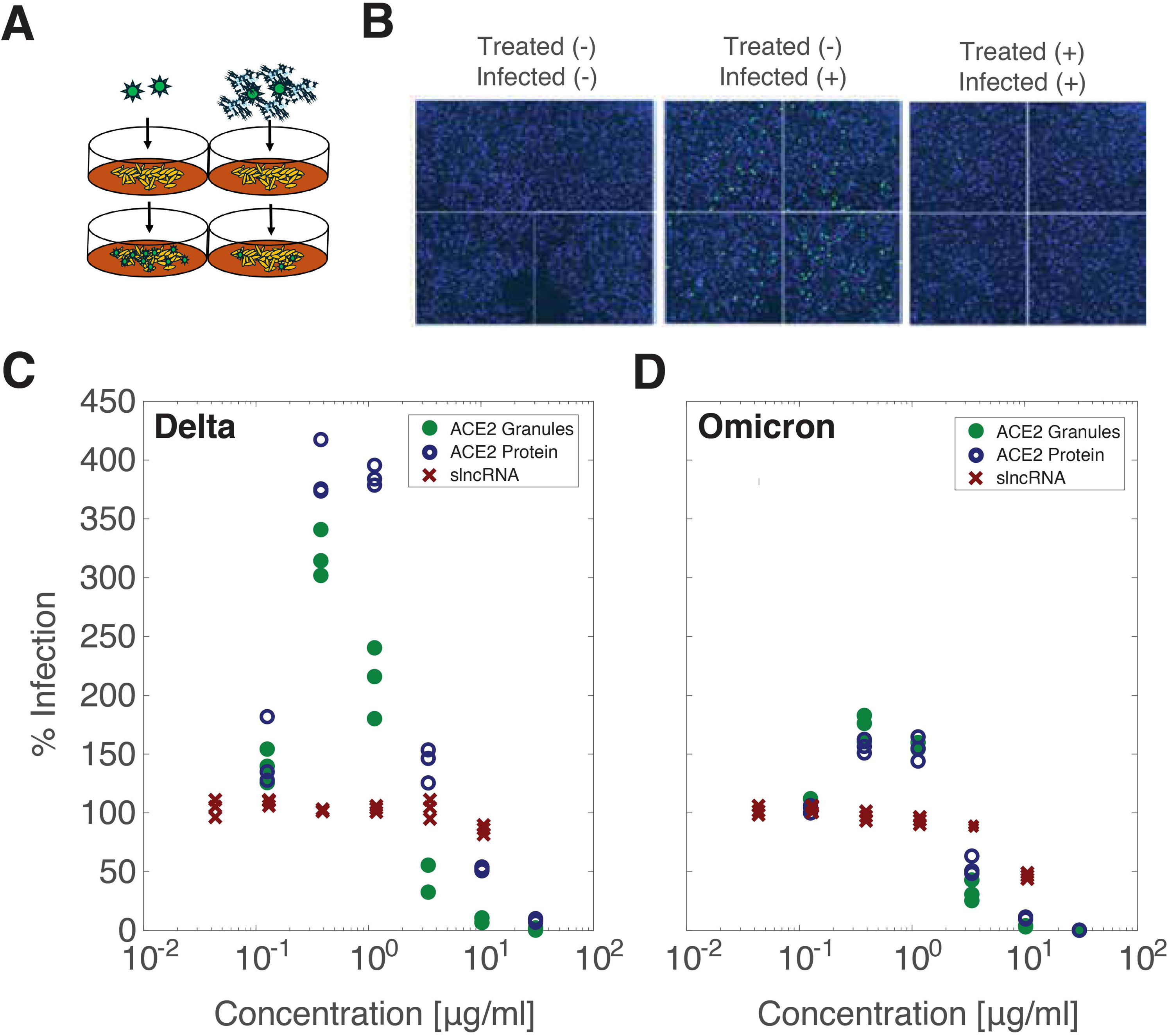
Interaction with ACE2 decoy particles enhances viral infection in low concentration and inhibits it at high concentrations. (**A**) schematic of viral entry assay. Virions (green objects) infected Vero E6 cells (yellow objects) in the absence (left) or presence (right) of decoy granules (**B**). Representative FoVs of infection by the Delta variant from the highest tested ACE2 decoy particle concentration. Blue signal – DAPI (cell). Green signal – AF488 (virus). **(C-D).** Normalized entry assay results for either the Delta (**C**) or Omicron (**D**) variants, in various ACE2 decoy particles concentrations. Below 3µg/ml, ACE2 decoy particles demonstrate enhancement of infection, while above viral infection is fully inhibited.

### Infection enhancement can be explained by spike priming

Given the infection enhancement effect observed for both Omicron and Delta for the protein alone and to a lesser extent for the granules, we wanted to develop a simplified kinetic model for infection in entry experiments that could provide a possible explanation for this unexpected result. Since a phase-separated environment (the decoy particles) might behave differently than a homogenous solution of soluble receptors, we first simulated a simplified infection process that we term “disrupted diffusion” (Figure 4A, top-left schema). We hypothesized that a diffusion process[26] controls the underlying dynamics of the virion search for its cellular target. Adding large decoy granules to the mix effectively introduces quasi-static obstacles positioned at random locations in space. Therefore, in this simplified scheme, the decoy granules serve as large static obstacles (RNA-only and sRNP granules measure ∼1 μm which is 10-fold bigger than a SARS-CoV-2 virion), where the smaller dynamic virions bind transiently and then bounce to the next decoy target in a stochastic pinball-like manner, leading to a disrupted diffusion process (Figure 4A, bottom-right schema). The simulation shows (Figure 4A, heatmap) that the disrupted diffusion leads to an increase in the average diffusion time per virion (Figure S4), which in turn translates to a reduction of the virion flux on the cell layer, resulting in inhibition of infection at high particle concentration, precisely as observed with the slncRNA-only sample (i.e. slncRNA-PCP-14x/MCP-15x phase separates on its own[26]).

**Figure 4-.**
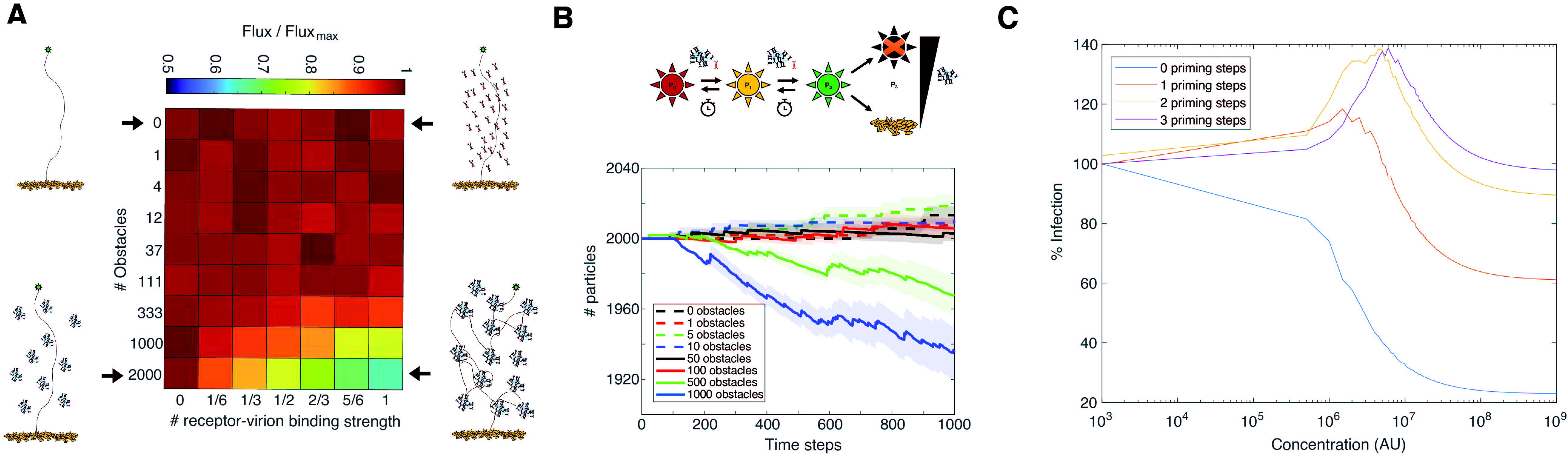
Model for infection in entry assays. (**A**). Normalized heatmap of virus infection flux (see Methods) in entry assays. Increasing the number of diffusion obstacles (y-axis) or the strength of their interactions (x-axis) results in decreased viral flux. Accompanying schemas: (top left) no obstacles. (Top right) Soluble decoy proteins. (Bottom left) Obstacles without a specific target. (Bottom right) Obstacles with high interaction specific target. (**B**) Priming model schema (top) and results (bottom). Prior to cell infection, a SARS-CoV-2 virion encounters either an ACE2 decoy particle or soluble ACE2, affecting one of its RBDs. A third priming event results in infection (if done in the cell layer) or virus elimination (if done by a decoy particle). Without decoys or free proteins, over time the priming is reversed. At low obstacle numbers (<100, a virus:obstacle ratio of 20:1), viral titer increases, while higher number of obstacles reduce the viral titer. (**C)** The model described in equations 5-10 was solved via MATLAB for multiple primed state with the following values for the constants: 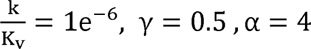.

The experimentally observed enhancement of infection at lower protein titer was not observed in our initial simulations using the simple disrupted diffusion model. In an attempt to provide a simple biological mechanism, we considered added “priming” steps to the disrupted diffusion simulation. We define priming as an effect that occurs prior to virus activation. In this case, at least one ACE2-spike binding event must occur at either an obstacle or the cell wall, for the virus to become activated, or primed, for infection at a subsequent interaction with the cell wall. Qualitatively, in the case of a low concentration of obstacles, a virion may encounter one or two obstacles before reaching the cell surface. This implies that while the flux of virions on the cell wall will somewhat decrease, as expected due to the presence of the obstacles, the actual flux of primed virions may increase over some low range of obstacle concentration, leading to enhancement of infection (Figure 4B). In addition, we simulated priming by setting up kinetic equations where the virus transitions from unprimed to as many as 3 primed states[35] in a stepwise fashion (see model section in Methods). The simulations for various number of priming steps (Figure 4C) show that at two priming steps and above, enhancement of infection occurs in a fashion that is similar to what was observed in the SARS-CoV-2 infection experiments. Interestingly, the model indicates that a higher number of priming steps leads to a stronger enhancement effect, suggesting that the difference between the antiviral response recorded for Omicron and Delta variants may stem from the latter requiring one more priming step than the former. Consequently, the inhibition patterns generated by the various antiviral decoy granules seem to reflect different underlying molecular and kinetic mechanisms associated with infection, which for the case of AntiCoV decoy leads to increased potential efficacy as compared with the decoy soluble proteins alone.

## Discussion

In this work, we build on our previous studies to show that phase-separated gel-like synthetic RNA-protein granules can be adapted to function as antiviral decoy nanoparticles that inhibit SARS-CoV-2 infection (Figure 3). Our particles remain stable at room temperature over several days (Figure 1) and retain functionality for at least several hours once exposed to either viral particles or components thereof (Figure 2-3). We further show that the mode of action is likely adsorption of the virions to the phase-separated particles, by demonstrating *in vitro* that the receptor binding domain of the spike protein is absorbed into the granules in a sponge-like fashion, forming stable filament-like structures composed of tightly bound slncRNA and proteins (Figure 2). In particular, using FRET we demonstrate an average separation of ∼6 to 8 nm between the fluorescent moieties of both ACE2 and the SARS-CoV-2 RBD, indicating a stably bound biomolecular complex spread throughout the granular structure (Figure 2). Finally, when either the granules or the stabilized ACE2 receptor were mixed with either the Delta or Omicron (BA.1) variants of SARS-CoV-2 in a viral entry assay, complete inhibition of infection of Vero E6 cells was observed (Figure 3). Consequently, both the stabilized soluble ACE2-mCherry-tdPCP receptor fusion protein and the phase-separated ACE2-slncRNA decoy granules can now be considered as pan-variant candidate prophylactic therapeutics that can potentially be used separately or in addition to vaccines to prevent SARS-CoV-2 infection.

The recent SARS-CoV-2 pandemic has highlighted limitations of our current pandemic control measures. First, global vaccination efforts during pandemic and non-pandemic times are limited by the need to distribute and/or store the vaccines in cold storage (i.e. −20 °C or lower). This implies that nearly half the global population cannot gain access to these vaccines[36–38]. Second, vaccines require administration by trained personnel and are typically delivered via intravenous injection. Third, vaccines need to be constantly redesigned for pandemic-potential RNA-based respiratory viruses to keep up with escape mutations that can render them obsolete within a few months, which adds a huge R&D and manufacturing burden to drug companies, and may require the population to get vaccinated on at least an annual basis. This, in turn, leads to reduced population compliance with vaccination guidelines, as is observed historically for the influenza virus vaccine and more recently for SARS-CoV-2. Our antiviral decoy granules have the potential to address all of these limitations insofar as preventing SARS-CoV-2 infection is concerned. The stability of the particles may facilitate an easy-to-deliver formulation (e.g. throat or nasal spray) that does not necessitate medical intervention and can be stored for long durations at room temperature or in refrigerated conditions (i.e. 2-8 °C). The decoy approach, which targets the evolutionary bottleneck of virion entry into the cell, ensures that a decoy-based therapeutic formulation will display strong pan-variant efficacy as was shown here for the Delta and Omicron variants. This means that a granule- or receptor-based therapeutic can be designed and prepared in advance to treat an emerging SARS pandemic, thus allowing health authorities to have an immediate countermeasure while new vaccines are being developed.

Finally, we note that while promising, our AntiCoV ACE2 decoy granules showed a complex dose response behavior to viral infection (Figure 3-4), suggesting a more nuanced receptor-RBD mode of interaction. While enhancement of infection by SARS2 variants at low ACE2 concentrations is a controversial subject and has only been observed in very few reported studies[39–41], the results shown here provide an impetus for further research into this phenomenon to confirm whether or not it is a real feature of the SARS-CoV-2 infection process or an artifact of the Vero assay used here and elsewhere[39,40]. In particular, the proposed mechanism of virus priming (Figure 4B-C) is consistent with several structural analysis studies that were carried out on ACE2-spike complexes[42–49]. Specifically, encounters between the spike protein and ACE2 induce changes in the RBD conformation (i.e. priming), and different variants maintain different basal levels of the open/up RBD conformations within the spike trimer (i.e. initial availability for binding). Therefore, while it is entirely possible that the simplified infection models devised here may only be relevant to the cell-types and entry assays used in our experiments, they may nevertheless shed light on complex kinetic phenomena that play a crucial role in SARS-CoV-2 infection. Consequently, our platform constitutes not only a new potentially broad-spectrum therapeutic but also provides an alternative assay for the discovery and study of viral infection processes, which could ultimately yield new antiviral therapeutic strategies.

### Concluding Remarks

This work demonstrates that phase-separated RNA-protein condensates can be engineered into stable, multivalent antiviral decoys that neutralize SARS-CoV-2 variants without the need to store them at low temperatures. By absorbing viral RBDs with nanometer-scale proximity to ACE2 and achieving full inhibition of Delta and Omicron entry in vitro, AntiCoV particles showcase a new strategy that leverages phase separation to immobilize and inactivate virions in ways conventional biologics cannot. The modularity of the slncRNA scaffold suggests potential for rapid reprogramming and even broad-spectrum prophylaxis if multiple receptors can be co-displayed. However, further developmental work remains (see Outstanding Questions), including how granule stability relates to structural organization, whether multiplexed receptors can function synergistically, and whether the low-dose enhancement seen *in vitro* might pose limitations *in vivo*. Addressing these challenges will determine how far this platform can advance toward practical antiviral deployment.

### Outstanding Questions

- **How are granule stability and antiviral function linked at the structural level?** The slncRNA cassettes show multi-week stability by qPCR, while microscopy reveals slowly evolving granule morphology and increasing particle volume over time. It remains unclear how structural features (e.g. hairpin packing, protein density, or phase-separation state) determine the granule’s ability to retain ACE2, resist degradation, and maintain antiviral potency. A deeper understanding of how nanoscale organization relates to functional longevity could guide rational engineering of more durable decoy systems.
- **Can multivalent condensates be expanded into broad-spectrum antiviral prophylactics?** In principle, the modularity of the tdPCP-slncRNA scaffold allows co-display of multiple recombinant receptors (e.g., ACE2 against coronaviruses, sialoproteins against influenza viruses, heparan-sulfate against respiratory syncytial virus, etc.). However, it remains unknown whether mixing distinct receptors within a shared condensate preserves proper orientation, affinity, accessibility, or competitive binding dynamics. Determining whether granules can be engineered to neutralize multiple viral families simultaneously is a key question for future development.
- **What drives the infection enhancement observed at low decoy concentrations?** Both soluble ACE2 and ACE2 granules show enhancement of infection in Vero entry assays at low doses. It is unclear whether this phenomenon reflects a cell-line artifact, a property unique to SARS-CoV-2 or its variants, or a more general priming-driven mechanism of viral entry. Establishing whether enhancement is conserved across cell types, viral strains and families, and receptor decoy designs will be crucial for predicting safety, optimizing dosing, and guiding regulatory considerations.

### Technology Readiness

The AntiCoV decoy platform is currently at an early preclinical stage, corresponding to Technology Readiness Level (TRL) 2-3. Our results demonstrate that phase-separated RNA-protein granules displaying ACE2 are structurally stable for extended periods at room temperature and can fully inhibit SARS-CoV-2 entry in vitro against both Delta and Omicron variants. These findings validate the fundamental technological concept and establish a reproducible in vitro performance benchmark. The biophysical characterization (confocal imaging, FRET) further supports the mechanistic feasibility of virus absorption into the granule matrix, while the kinetic model clarifies the biphasic dose response observed in entry assays. Together, these results indicate a functional proof of principle with clear potential for extension to additional viral receptors.

Despite these encouraging advances, several challenges must be addressed before progressing to translational development. First, the granules must be evaluated in physiologically relevant environments, including mucosal fluids, primary airway cells, and animal models, to assess stability, biocompatibility, and *in vivo* antiviral efficacy. Second, scalable and GMP-compatible manufacturing processes for both the RNA scaffolds and protein components will need to be optimized, including purification, formulation, and long-term storage. Third, the immunogenicity of repeated exposure to synthetic RNA-protein condensates must be evaluated carefully, as well as their clearance pathways and potential interactions with endogenous biomolecular condensates. Regulatory considerations will also influence deployment strategies, particularly for prophylactic use in healthy individuals. If these challenges can be addressed, AntiCoV decoy particles could represent a low-cost, cold-chain-independent antiviral technology suitable for rapid deployment during future respiratory virus outbreaks. More broadly, the platform may enable a new class of modular, phase-separated antiviral decoys targeting diverse viral entry receptors.

## Supporting information

Supplemental Figures and Sequences

## Acknowledgements

We thank the Life Sciences and Engineering (LS&E) Infrastructure Center at the Technion (Dr. Nitsan Dahan and Dr. Yael Lupu-Haber from the microscopy core facility) for their excellent technical assistance with confocal microscopy imaging. We would like to thank Viroclinics Inc. for executing the Vero E6 viral entry assay in GLP conditions. This work was greatly supported by the European Union’s Horizon 2020 Research and Innovation Programme under grant agreements no. 851615 and no. 851065.

## Author Contributions

**OW** designed and carried out experiments and analysis for most of the data together with **TS**. **NG** and **RA** conceived the priming model. **NG** coded the priming model and the virus infection simulations and inferred the protein structures using AlphaFold. **OW** and **TS** performed the slncRNA synthesis and qPCR assays. **OW** and **SG** designed the plasmid for ACE2 and **OW** designed and cloned its mutant variants. **SG** guided the microscopy experiments and image analysis. **RA** supervised the study. **OW** and **RA** wrote the manuscript.

## Declaration of Interests

The authors declare the following competing financial interest(s): **OW**, **NG**, **SG**, and **RA** are inventors on US Provisional Patent Application No. 63/187969 concerning some of the technologies described. **RA** is an inventor on US Patent Application 2021/0095296 A1.

## STAR⍰Methods

### Key resources table

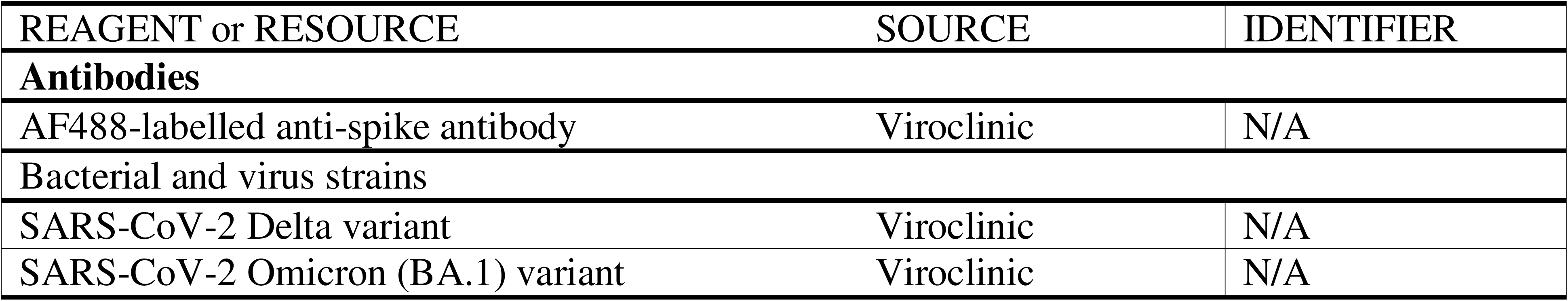
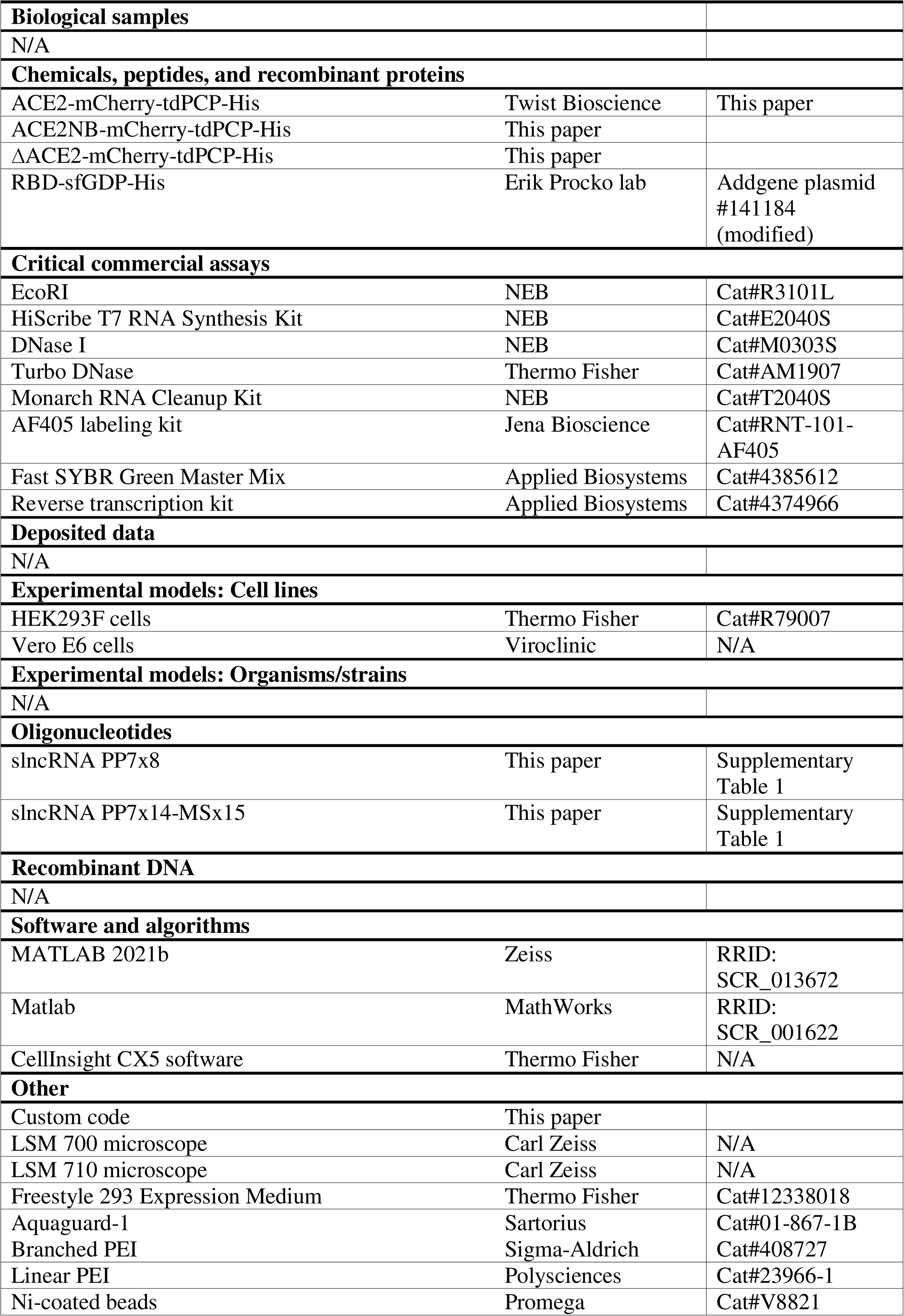
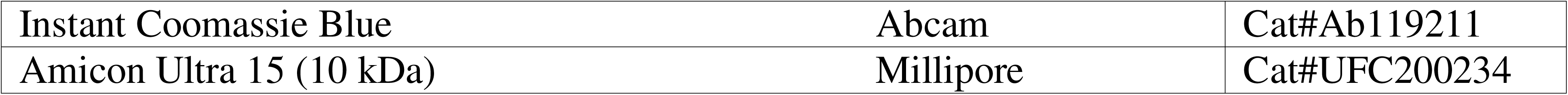

### Lead contact

Further information and requests for resources and reagents should be directed to the lead contact.

### Materials availability

All plasmids and materials generated in this study are available upon request.

### Data and code availability

The numerical simulations and priming model were implemented in MATLAB 2021b. Code and instructions for use are available at 10.5281/zenodo.17660183.

### Plasmid construction

All proteins were expressed in female HEK293F cells (Thermo Fisher cat. R79007) and purified using a C-terminus affinity tag comprised of six histidines (his-tag), positioned immediately upstream to the stop codon of all coding sequences. The sequence of human ACE2, followed by an mCherry-tdPCP-his component, was cloned by Twist Bioscience into the pTwist-betaglobin construct. The ectodomain of ACE2 was extracted from UniProt and contained the protein’s native signal peptide and the extracellular domain, namely amino acids 1-740. We also designed the following mutants: a non-RBD-binding mutated variant (ACE2NB, a D355A mutation[30]) and a non-dimerizing ACE2 (ΔACE2, a deletion of amino acids 616-726[31]). pcDNA3-SARS-CoV-2-S-RBD-sfGFP was a gift from Erik Procko (Addgene plasmid #141184), and a his-tag was added to the C-terminus of the RBD-sfGFP open reading frame (ORF). Plasmids containing the synthetic long non-coding RNA (slncRNA) cassettes were created as previously described[26]. All sequences can be found in Supplementary Table 1.

### In vitro transcription and purification of RNA cassettes

RNA transcription of synthetic RNA cassettes was done as previously described[26]. Briefly, a vector containing the slncRNA DNA sequence, flanked by two EcoRI restriction sites, was digested with EcoRI-HF (NEB, cat. R3101L) per the manufacturer’s instructions to form a linear fragment. The enzyme was then heat-inactivated by incubating the restriction reaction at 65 °C for 20 min. For fluorescently labeled RNA, 1µg of the restriction product was used as template for in vitro transcription using HighYield T7 Alexafluor-405 (AF405) RNA labeling kit (Jena Bioscience, cat. RNT-101-AF405), according to the manufacturer’s instructions. Non-fluorescent RNA was transcribed using the HiScribe™ T7 High Yield RNA Synthesis Kit (NEB, cat. E2040S). Following in vitro transcription by either kit, the reaction was diluted to 90 µl and was supplemented with 10 µl DNAse I buffer and 2µl DNase I enzyme (NEB, cat. M0303S) and incubated for 15 min at 37 °C to degrade the DNA template. RNA products were purified using Monarch RNA Cleanup Kit (NEB, cat T2040S) and stored in −80°C. For qPCR, RNA products were purified using Monarch RNA Cleanup Kit (NEB, cat T2040S) and stored in −80°C. RNA samples used in qPCR were subjected to Turbo DNase (Thermo Fisher, cat. AM1907) treatments according to manufacturer’s protocol, twice for 30min, and subsequently incubated with DNase inhibitor reagent of the same kit according to manufacturer’s instructions.

### HEK293F cells maintenance

A cryotube with 1ml with 10%v/v DMSO (Sigma-Aldrich, cat. D2650) of 10^7^ HEK293F cells was thawed as follows: the cryotube’s content was transferred into 4ml fresh Freestyle media (Thermo Fisher, cat. 12338018), prewarmed to 37°C, and then immediately centrifuged at 500xg for 5min. Supernatant was removed, and cell pellet was resuspended i 7ml of fresh Freestyle media and transferred to 23ml of prewarmed Freestyle media in a 125ml flat-bottom flask (TriForest, cat. TF FPC0125S). Cells were then moved to an incubator and grown under 37°C, 8% CO_2_ and humid conditions (an in-hood basin with 500ml autoclaved DI water, supplemented with 1%v/v Aquaguard-1 solution (Sartorius, cat. 01-867-1B)). The cells were placed on an in-incubator orbital shaker which rotated at 135rpm. These conditions were in accordance with the cells’ manual.

The cell concentration exceeded 1.2×10^6^ cells/ml 3-4 days after thawing, at which point cells were passed by diluting them 1:6 in a final volume of 30ml fresh growth media. Cells were routinely passed every 3-4 days, as cell concentration was sufficient. Cells were checked for mycoplasma in accordance with manufacturer protocol (Vazyme, cat. D101-02).

### Cell transfection

24h before transfection, HEK239F cells were seeded at 0.6-0.7×10^6^ cells/ml in 30ml fresh Freestyle media and allowed to grow overnight. On the day of transfection, cells were diluted to 1×10^6^ cells/ml. Per flask containing 30ml of culture, up to 40µg of plasmid DNA was diluted in OptiMEM buffer (Thermo Fisher, cat. 31985070) to a final volume of 600µl. 120µl of 0.5mg/ml branched PEI solution (Sigma-Aldrich, cat. 408727) or 60µl of 1mg/ml linear PEI (Polysciences, cat. 23966-1) was diluted in OptiMEM buffer to a final volume of 600µl. The PEI-OptiMEM solution was added to the DNA-OptiMEM solution (final mixed volume of 1.2ml) and incubated at room temperature for 15min. Each flask was then transfected with the appropriate plasmid within 5 minutes. Transfected cells were maintained for 5-7 days, during which time the growth media changed color to light red due to the mCherry-label in the recombinant proteins.

### Protein extraction from HEK293F cells

1ml of nickel (Ni)-coated beads (Promega, cat. V8821) were transferred into 50ml conical tubes and were allowed to settle. 400µl of supernatant was removed and beads were washed in 2.5ml of equilibration buffer (50mM NaH_2_PO_4_, 300mM NaCl, 5mM imidazole, pH 8). After the beads resettled, 2ml of the supernatant was removed. Transfected cells were centrifuged 5-7 days post transfection at 5000rpm for 20min at 4°C. Up to 40 ml supernatant was transferred to a 50ml conical tube with settled beads while cell pellet was discarded. Bead-protein mix was incubated at room temperature for 1h in an end-over-end shaker. After incubation, the solution was transferred to a gravity column (Bio-rad, cat. 731150) and ∼100µl of the flowthrough was collected. Next, beads were washed three times using 5ml of wash solution (50mM NaH_2_PO_4_, 300mM NaCl, 20mM imidazole, pH 8), and ∼100µl was collected from the first wash. Finally, beads were resuspended inside the column in 2ml elution buffer (50mM NaH_2_PO_4_, 300mM NaCl, 500mM imidazole, pH 8) and left to incubate for 15min, after which the entire elution volume was collected in 2-3 fractions.

### Verification of extracted proteins using SDS-PAGE

15µl of each fraction (flowthrough, wash, elution) per protein was taken to sodium dodecyl sulfate polyacrylamide gel electrophoresis (denaturing SDS-PAGE), using a 10% acrylamide concentration in the resolving gel. 0.1mg/ml of purified BSA (NEB, cat. B9000) was used as positive control. Samples were boiled at 95°C for 10min before loading onto the gel. Gel running conditions were 150V for 70min. Afterwards, the gel was stained using Instant Coomassie Blue (Abcam, cat. Ab119211) for 15min.

### Buffer exchange of extracted proteins

The high imidazole content from the elution buffer was diluted ∼1:10^4^ by using an Amicon filtration tube (Millipore, cat. UFC200234) according to the manufacturer’s protocol. In brief, PBSx10 was added to the eluted protein samples as 1/9 volume equivalent its post-extraction volume. Then, PBSx1 was added to dilute the imidazole concentration to 100mM, the maximal concentration recommended by the Millipore protocol. The diluted protein solution was loaded onto the Amicon tube and centrifuged for 15min at 5000rpm several times, until reaching 400-500µl retentate volume. In between each centrifuge, protein precipitate was resuspended by carefully pipetting inside the column. Once arriving at the appropriate volume range, 500µl of PBSx1 was added to the protein solution inside the column, diluting the imidazole ∼2-fold. This process was repeated eleven times.

### Formation of RNA-protein granules

For FRET assays, 250fmol slncRNA (PP7×8 cassette) was mixed with each of the proteins at a molar ratio of 100:1 protein-to-RNA, using a 7µl of granule buffer (750mM NaCl, 1mM MgCl_2_, 10% PEG 4000 final concentrations) and brought with water to a final volume of 21µl. For confocal stability assays and viral entry assays, 10:1 protein-to-slncRNA ratio was used (2.5pmol of PP7×8 slncRNA cassette and 10pmol of PP7×14-MSx15 slncRNA cassette, respectively). Stability assays maintained a final volume of 21µl, while for viral assays, a final volume of 210µl was used for a four-replicate mix (52.5µl for each replicate, see viral entry assay method protocol). In each case, the slncRNA-protein mixture was incubated at room temperature for 1h before being taken for imaging. For the viral assay, ACE2 decoys or their moieties were shipped on ice and frozen upon arrival before use.

### Quantitative PCR (qPCR)

Stock tube of 3.75pmol of slncRNA was incubated in granule buffer for 1h at room temperature, in a final volume of 315µl (completed with nuclease-free water). Samples were prepared as follows: 1.7µl of granules were then diluted 1:10 and was subjected to cDNA synthesis (Applied Biosystems, cat. 4374966), according to manufacturer’s protocol, resulting in a further 1:2 dilution. As slncRNA control, a similar, non-reverse-transcribed granule sample was diluted 1:20, matching the final granule concentration post-cDNA synthesis. After cDNA synthesis was completed, cDNA and slncRNA control were amplified in qPCR, using dedicated primers and fast SYBR (Applied Biosystems, cat. 4385612), where the sample cDNA or RNA comprised 40% of the reaction (final volume 10µl). Samples were loaded in triplicates. Importantly, samples without the 1:10 dilution step prior to cDNA synthesis were not successfully amplified.

After the 1h period of incubation on Day 0, 58.8µl of granules from the stock tube were aliquoted in LoBind tubes, and stored at either −80°C, −20°C, 4°C, room temperature (RT), and 37°C. On days 4 and 7, samples were left for 30min at room temperature for thermal equilibrium before being prepared for cDNA synthesis.

### Confocal microscopy imaging

For all imaging, we used the LSM-700 (Carl Zeiss) confocal microscope, equipped with a BIG (GaAsP detector) unit, using a Plan-Apochromat 63x/1.40 Oil objective. A pinhole of 1.5µm was set for each laser. Emission windows were 578-800, 300-583, and 300-483 for the 555nm (mCherry), and 405nm (AF405) channels, respectively, for all assays. For each assay we used standard microscopy slides with 1.5H coverslips, and each sample was imaged using 5µl.

Granules or their protein or slncRNA moieties were incubated and maintained at room temperature for a week and imaged on days 0, 4, and 7 post-mixing of proteins and slncRNA (Day 0 imaging took place 1h post-mixing). Granules and moieties were kept in room temperature, covered with aluminum foil and stored in a drawer to protect from light.

### Viral infection of Vero E6 Cells with SARS-CoV-2 and analysis

The experiments were carried out in a GLP-certified dedicated EU BSL3 (Bio-Safety Level 3) facility (see Acknowledgements). Female Vero E6 cells[32] were seeded in a 96-wells plate (columns 1 to 11) in 100µl of M199 growth media (Gibco, cat. 41150087, supplemented with 5%v/v FBS) and allowed to adhere and grow overnight for 90% confluency (culture plate). The next day, a parallel plate was prepared (infection plate), holding both ACE2 granules and coronavirus virions (Delta or Omicron BA.1 variants). ACE2 and RNA highest tested concentration was concentrations were ∼0.2µM and ∼0.02M, respectively. All tested conditions were diluted 3-fold seven times in triplicates. The plate was prepared as follows: 52.5µl of ACE2 granules, or ACE2-only or RNA-only were added to row A of columns 1-3, 4-6, and 7-9, respectively. 52.5µl of infection media (M199, supplemented with 0.3% BSA) was added to columns 10-12, where untreated infected cells, untreated uninfected cells, and no cells controls would populate columns 10, 11, and 12, respectively. To rows B through H, 35µl of infection media was added. Using multichannel pipette, 17.5µl of the content of row B was transferred to row C, and the process repeated until row G. This comprised dilutions 0-7. 35µl of virus (∼2.5×10^5^ PFU/ml) was added to each well except uninfected columns (columns 11 and 12) which received 35µl of infection media instead. Each well in the infection plate then held 70µl of infection media-virus-test article. The infection plate was incubated for 1h at 37°C in a humidifier incubator. Following the incubation time, media was replaced with fresh media in the culture plate and the infection plate content was transferred to the culture plate. Infection was allowed to take place for 6h inside the incubator. Cell density was ∼2×10^4^ cells/well, with an MOI of ∼0.5.

Following the infection period, media was removed, and cells were washed once with PBS. Then 100µl of 4% formaldehyde in PBS was added to each well for fixation of cells. Plates were incubated at room temperature protected from light for 30min, and afterwards washed once with PBS. Cells were stored in 100µl fresh PBS at 4°C and stained within 2 days using AF488-labelled anti-spike antibody. Subsequently, plates were analyzed using CellInsight CX5 (Thermo Fisher).

### Numeric model for virion flux measurement

We developed a simulation model to study a disrupted diffusion process of particles in a 3D bounded space with obstacles, incorporating particle attachment, hit counts, and particle generation mechanisms. The primary goal was to analyze the flux of particles hitting a cell layer, i.e., “the measurement wall”, arbitrarily designated as the x = 0 plane.

Model assumptions include (1) 3D bounded space with limits at [*L_X_,L_Y_,L_Z_*]; (2) particles undergo Brownian motion, characterized by a diffusion coefficient *D*; (3) virion flux is measured on the plane of *x*= 0; (4) the space contains *K* spherical obstacles which simulate a phase-separated condensate containing decoy receptors; and (5) upon hitting an obstacle, a virion becomes immobilized for a specified number of time steps *t_attach_* before being released in a random direction.

The overall number of virion particles remained the same, and whenever a virion reached x = 0, a new virion was generated randomly in the 3D space. At the start of the simulation, *N* particles and *K* obstacles are initialized at random positions withing the bounded space. At each simulation step, particle positions are updated based on a diffusion process (equation #2):

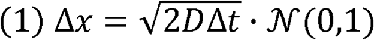

Where *N*(0,1) represents a number randomly generated from a standard normal distribution. Particles are reflected back into the bounded space if they hit a boundary, as follows (equation #3):

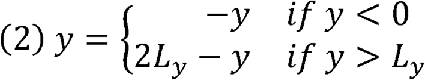

Similar reflection is applied for the *z* dimension.

For the *x* dimension the following applies (equation #4):

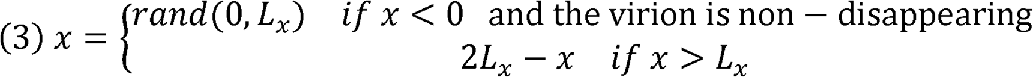

Where *rand*(0, *L_X_*) indicates a number randomly generated from a uniform distribution in the range [0,*L_X_*].

The number of particles hitting the measurement wall is recorded, and the flux rate is calculated at the end of the simulation as (equation #5), where *S* is the total number of simulation steps (10,000):

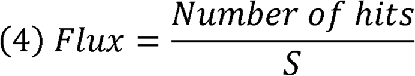

### Priming model

In order to model virus priming in a controlled cell culture setting, we have set up the following system of equations for *n* priming steps:

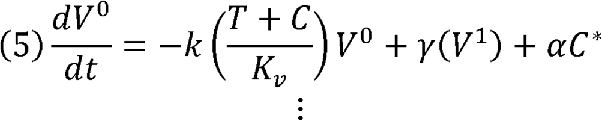

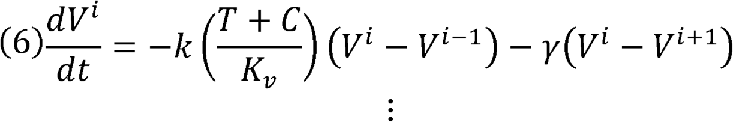

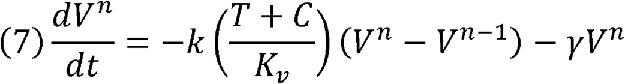

Where *V^0^*, *V^i^*,and *V^n^*correspond to the unprimed, i^th^-step, and n^th^-step primed virus concentration respectively. *T* and *C* correspond to the therapeutic and uninfected cell concentrations respectively. *C\** corresponds to infected cell concentration which creates new unprimed virus particles at a rate α. *K_v_*is the virus binding affinity to ACE2 (either as a therapeutic or on the cell), *k* is the rate at which the virus binds ACE2 leading to an increase in the “primed” state of the virion, and γ corresponds to a spontaneous reversion of the virion to a lower “primed” state. The priming equations are then complemented by the following equations:

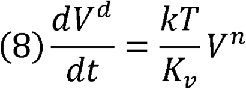

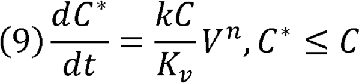

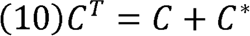

where *V^d^* corresponds to the concentration of deactivated virus particles as a result of the binding of a fully primed virion to a decoy therapeutic, the rate of creation of infected cells *C\** is provided by the probability for a fully primed virion (*V^n^*) to interact with an uninfected cell (*C*), and d *C^T^* is a constant corresponding to the total cell concentration in the simulation (1e6 in our settings). Simulation was implemented in Matlab 2021b (The MathWorks). See code availability statement for more detail.

## References

1. Magden, J. et al. (2005) Inhibitors of virus replication: recent developments and prospects. Appl Microbiol Biotechnol 66, 612–621

2. Marzi, M., et al. (2022) Paxlovid: Mechanism of Action, Synthesis, and In Silico Study. BioMed Research International 2022, 7341493

3. Bakheit, A.H. et al. (2023) Chapter Three - Remdesivir. In Profiles of Drug Substances, Excipients and Related Methodology 48 (Al-Majed, A. A., ed), pp. 71–108, Academic Press

4. Singh, A.K., et al. (2021) Molnupiravir in COVID-19: A systematic review of literature. Diabetes & Metabolic Syndrome: Clinical Research & Reviews 15, 102329

5. Jordheim, L.P. et al. (2013) Advances in the development of nucleoside and nucleotide analogues for cancer and viral diseases. Nat Rev Drug Discov 12, 447–464

6. Huang, D. et al. (2021) Efficacy and safety of umifenovir for coronavirus disease 2019 (COVID-19): A systematic review and meta-analysis. Journal of Medical Virology 93, 481–490

7. Gupta, Y.K. et al. (2015) The Tamiflu fiasco and lessons learnt. Indian Journal of Pharmacology 47, 11

8. Jang, W.D. et al. (2021) Drugs repurposed for COVID-19 by virtual screening of 6,218 drugs and cell-based assay. Proceedings of the National Academy of Sciences 118, e2024302118

9. Riva, L. et al. (2020) Discovery of SARS-CoV-2 antiviral drugs through large-scale compound repurposing. Nature 586, 113–119

10. Henderson, D.A. (2011) The eradication of smallpox – An overview of the past, present, and future. Vaccine 29, D7–D9

11. Holzmann, H. et al. (2016) Eradication of measles: remaining challenges. Med Microbiol Immunol 205, 201–208

12. Bandyopadhyay, A.S. et al. (2015) Polio Vaccination: Past, Present and Future. Future Microbiology 10, 791–808

13. James, E.R. (2021) Disrupting vaccine logistics. Int Health 13, 211–214

14. Bollyky, T.J. et al. (2020) The Equitable Distribution of COVID-19 Therapeutics and Vaccines. JAMA 323, 2462–2463

15. Md Khairi, L.N.H., et al. (2022) The Race for Global Equitable Access to COVID-19 Vaccines. Vaccines 10, 1306

16. Starr, T.N. et al. (2021) Prospective mapping of viral mutations that escape antibodies used to treat COVID-19. Science 371, 850–854

17. Chan, K.K. et al. (2020) Engineering human ACE2 to optimize binding to the spike protein of SARS coronavirus 2. Science 369, 1261–1265

18. Leach, A. et al. (2021) A tetrameric ACE2 protein broadly neutralizes SARS-CoV-2 spike variants of concern with elevated potency. Antiviral Res 194, 105147

19. Torchia, J.A. et al. (2022) Optimized ACE2 decoys neutralize antibody-resistant SARS-CoV-2 variants through functional receptor mimicry and treat infection in vivo. Science Advances 8, eabq6527

20. Guo, L. et al. (2021) Engineered trimeric ACE2 binds viral spike protein and locks it in “Three-up” conformation to potently inhibit SARS-CoV-2 infection. Cell Res 31, 98–100

21. Pinkert, S. et al. (2016) Soluble coxsackie- and adenovirus receptor (sCAR-Fc); a highly efficient compound against laboratory and clinical strains of coxsackie-B-virus. Antiviral Res 136, 1–8

22. Haim, H. et al. (2009) Soluble CD4 and CD4-Mimetic Compounds Inhibit HIV-1 Infection by Induction of a Short-Lived Activated State. PLoS Pathog 5, e1000360

23. Li, G. et al. (2023) Engineered soluble ACE2 receptor: Responding to change with change. Front Immunol 13, 1084331

24. Blundell, P.A. et al. (2019) Insertion of N-Terminal Hinge Glycosylation Enhances Interactions of the Fc Region of Human IgG1 Monomers with Glycan-Dependent Receptors and Blocks Hemagglutination by the Influenza Virus. J Immunol 202, 1595–1611

25. Blundell, P.A. et al. (2020) Choice of Host Cell Line Is Essential for the Functional Glycosylation of the Fc Region of Human IgG1 Inhibitors of Influenza B Viruses. J Immunol 204, 1022–1034

26. Granik, N. et al. (2022) Formation of synthetic RNA protein granules using engineered phage-coat-protein −RNA complexes. Nat Commun 13, 6811

27. Granik, N. et al. (2025) Formation of Polyphasic RNP Granules by Intrinsically Disordered Qβ Coat Proteins and Hairpin-Containing RNA. ACS Synth Biol 14, 2081–2093

28. Katz, N. et al. (2021) Overcoming the design, build, test bottleneck for synthesis of nonrepetitive protein-RNA cassettes. Nat Commun 12, 1576

29. Kikuchi, N. et al. (2022) A Cell-Free Assay for Rapid Screening of Inhibitors of hACE2-Receptor-SARS-CoV-2-Spike Binding. ACS Synth Biol 11, 1389–1396

30. Brown, E.E.F. et al. (2021) Characterization of Critical Determinants of ACE2–SARS CoV-2 RBD Interaction. International Journal of Molecular Sciences 22, 2268

31. Yan, R. et al. (2020) Structural basis for the recognition of SARS-CoV-2 by full-length human ACE2. Science 367, 1444–1448

32. Ogando, N.S. et al. (2020) SARS-coronavirus-2 replication in Vero E6 cells: replication kinetics, rapid adaptation and cytopathology. J Gen Virol 101, 925–940

33. Chan, E.W. et al. (1980) Effects of polyethylene glycol on reverse transcriptase and other polymerase activities. Biochimica et Biophysica Acta (BBA) - Nucleic Acids and Protein Synthesis 606, 353–361

34. Bagnoli, J.W. et al. (2018) Sensitive and powerful single-cell RNA sequencing using mcSCRB-seq. Nat Commun 9, 2937

35. Fuentes-Prior, P. (2021) Priming of SARS-CoV-2 S protein by several membrane-bound serine proteinases could explain enhanced viral infectivity and systemic COVID-19 infection. Journal of Biological Chemistry 296

36. Fahrni, M.L. et al. (2022) Management of COVID-19 vaccines cold chain logistics: a scoping review. J Pharm Policy Pract 15, 16

37. Kis, Z. (2022) Stability Modelling of mRNA Vaccine Quality Based on Temperature Monitoring throughout the Distribution Chain. Pharmaceutics 14, 430

38. Mesa-Vieira, C. et al. (2021) Reprint of: The Dark Side of the Moon: Global challenges in the distribution of vaccines and implementation of vaccination plans against COVID-19. Maturitas 150, 61–63

39. Liu, Y. et al. (2021) An infectivity-enhancing site on the SARS-CoV-2 spike protein targeted by antibodies. Cell 184, 3452–3466.e18

40. Li, D. et al. (2021) In vitro and in vivo functions of SARS-CoV-2 infection-enhancing and neutralizing antibodies. Cell 184, 4203–4219.e32

41. Lusiany, T. et al. (2023) Enhancement of SARS-CoV-2 Infection via Crosslinking of Adjacent Spike Proteins by N-Terminal Domain-Targeting Antibodies. Viruses 15, 2421

42. Valério, M. et al. (2022) SARS-CoV-2 variants impact RBD conformational dynamics and ACE2 accessibility. Front Med Technol 4, 1009451

43. Pang, Y.T. et al. (2022) SARS-CoV-2 spike opening dynamics and energetics reveal the individual roles of glycans and their collective impact. Commun Biol 5, 1–11

44. Gur, M. et al. (2020) Conformational transition of SARS-CoV-2 spike glycoprotein between its closed and open states. J Chem Phys 153, 075101

45. Xu, C. et al. (2021) Conformational dynamics of SARS-CoV-2 trimeric spike glycoprotein in complex with receptor ACE2 revealed by cryo-EM. Science Advances 7, eabe5575

46. Cai, Y. et al. (2020) Distinct conformational states of SARS-CoV-2 spike protein. Science 369, 1586–1592

47. Lee, M. et al. (2023) Distinct Conformations of SARS-CoV-2 Omicron Spike Protein and Its Interaction with ACE2 and Antibody. Int J Mol Sci 24, 3774

48. Hong, Q. et al. (2022) Molecular basis of receptor binding and antibody neutralization of Omicron. Nature 604, 546–552

49. Zhao, Z. et al. (2022) Omicron SARS-CoV-2 mutations stabilize spike up-RBD conformation and lead to a non-RBM-binding monoclonal antibody escape. Nat Commun 13, 4958

