## Supplemental Figures and Sequences for "Synthetic RNA-protein decoy granules to prevent SARS-CoV-2 infection"

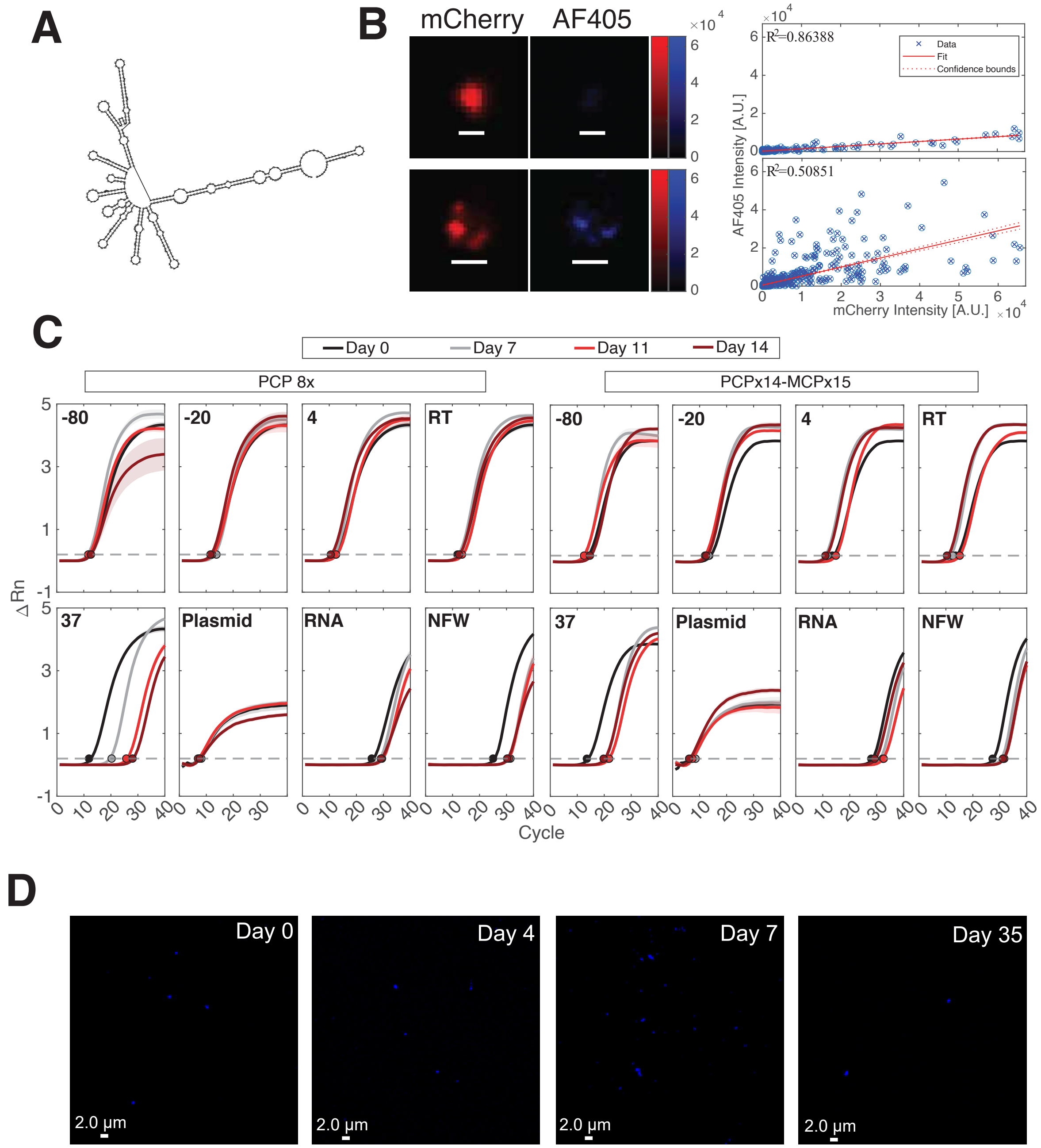


**Figure S1: Characterization of decoy particles stability, Related to Figure 1. (A)** Model structure computed using Vienna of a 478 nt slncRNA encoding 8 non-sequence repeating PCP hairpins. **(B)** (Left) Channels view of enlarged ACE2 decoy granules in Fig. 1D. (Right) Correlation curves between mCherry- and AF405-positive pixels of the two representative events, demonstrating variability between colocalization appearance and mathematical values. **(C)** qPCR data for the slncRNA PCPx14-MCPx15 cassette, used to generate Fig. 1F. Each hill curve represents the mean of three replicates, and shading area denotes standard error between replicates. Dashed line – threshold of 0.2. RNA concentration corresponds to a 1:20 diluted room temperature (RT) reaction (see Methods). NFW – nuclease free water. Plasmid concentration is 10ng/µl. D. Representative images of slncRNA-only (PCP-8x) over the course of 35 days, showing no meaningful change of size.

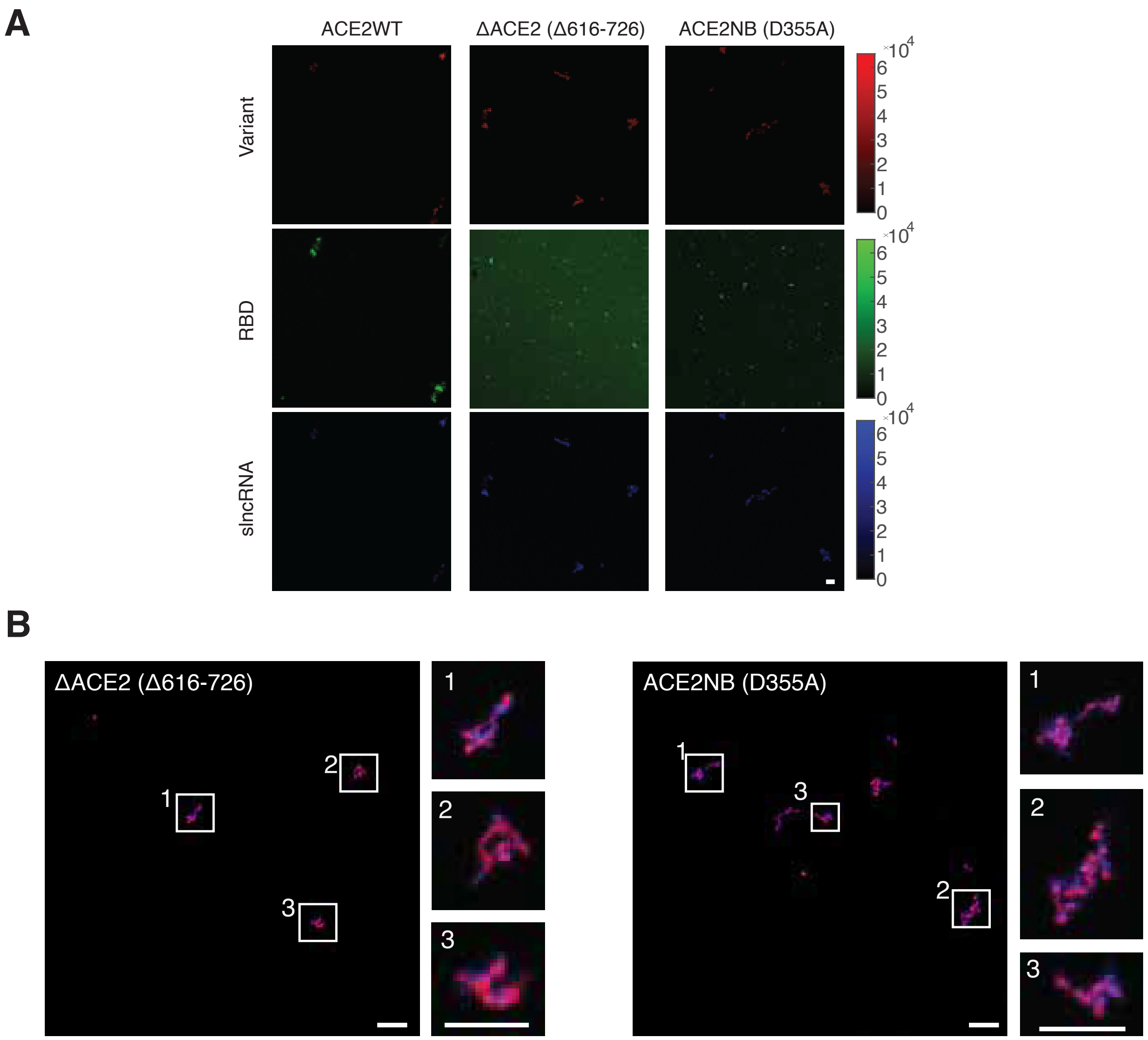


**Figure S2: ACE2 (wild-type and mutants) interactions with slncRNA and RBD, Related to Figure 2.**

**(A)** Individual channels of the ACE2 (wildtype and mutants) granules and RBD shown in Figure 2B, 2F-top left, and 2G-top left. **(B)** Granules formed by the ACE2 mutants.

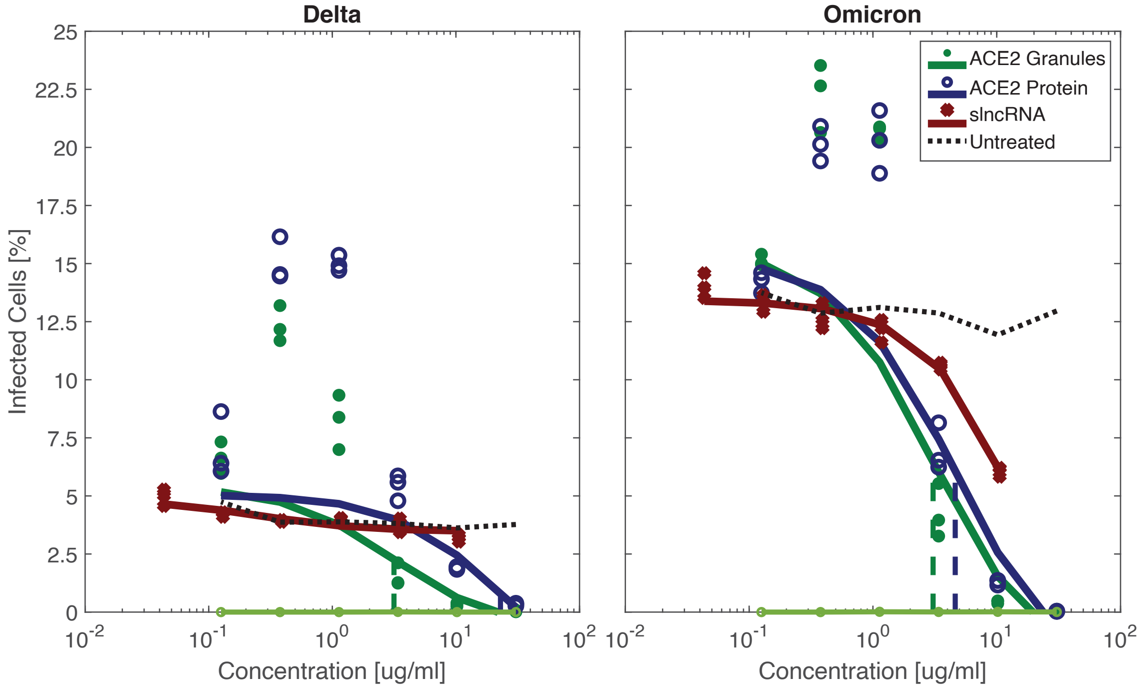


**Figure S3: ACE2 decoy particles inhibit SARS2 infection at a lower $IC_{50}$ value than an ACE2-only solution, Related to Figure 3.**

Infection by either the Delta (left) or Omicron (right) variants was inhibited in Vero E6 cells using ACE2 decoy particles. This was achieved at a lower $IC_{50}$ value for both variants.

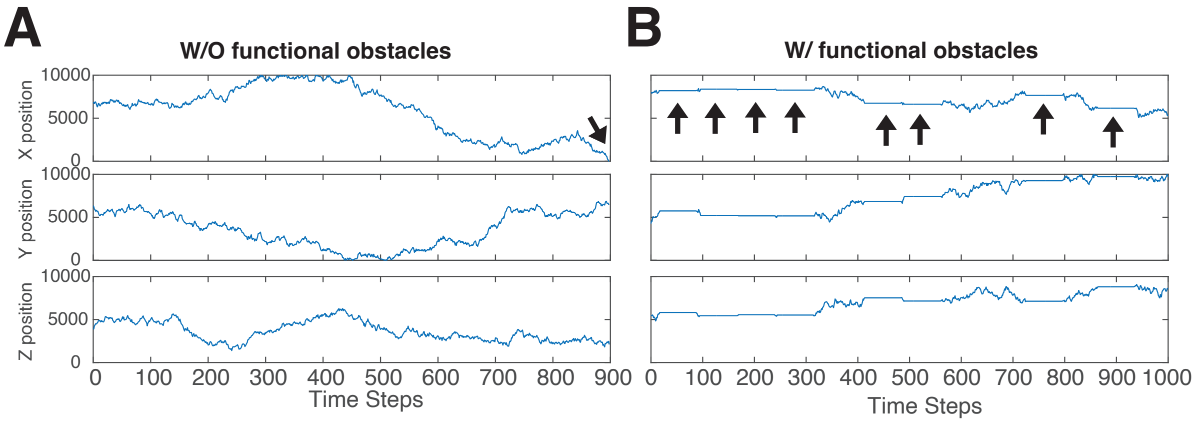


**Figure S4: Obstacles that interact with viral particles prolong viral diffusion, Related to Figure 4.**

**(A-B)** Representative trajectories in 3D space of a viral particle in the absence (A) or presence (B) of functional obstacles that it can interact with (i.e. ACE2 decoy particles). Interactions with an obstacle (marked by arrows in B) result in a delay in progression towards the cell layer (marked by an arrow in A, X position = 0) and increase the time it takes to infect a cell.

| Component | Sequence | Species | Molecular weight (kda) |
|---|---|---|---|
| ACE2-mCherry-tdPCP-His | MSSSSWLLLSLVAVTAAQSTIEEQAKTFLDKFNHEAEDLFYQSSLASWNYNTNITEENVQNMNNAGDKWSAFLKEQSTLAQMYPLQEIQNLTVKLQLQALQQNGSSVLSEDKSKRLNTILNTMSTIYSTGKVCNPDNPQECLLLEPGLNEIMANSLDYNERLWAWESWRSEVGKQLRPLYEEYVVLKNEMARANHYEDYGDYWRGDYEVNGVDGYDYSRGQLIEDVEHTFEEIKPLYEHLHAYVRAKLMNAYPSYISPIGCLPAHLLGDMWGRFWTNLYSLTVPFGQKPNIDVTDAMVDQAWDAQRIFKEAEKFFVSVGLPNMTQGFWENSLLTDPGNVQKAVCHPTAWDLGKGDFRILMCTKVTMDDFLTAHHEMGHIQYDMAYAAQPFLLRNGANEGFHEAVGEIMSLSAATPKHLKSIGLLSPDFQEDNETEINFLLKQALTIVGTLPFTYMLEKWRWMVFKGEIPKDQWMKKWWEMKREIVGVVEPVPHDETYCDPASLFHVSNDYSFIRYYTRTLYQFQFQEALCQAAKHEGPLHKCDISNSTEAGQKLFNMLRLGKSEPWTLALENVVGAKNMNVRPLLNYFEPLFTWLKDQNKNSFVGWSTDWSPYADQSIKVRISLKSALGDKAYEWNDNEMYLFRSSVAYAMRQYFLKVKNQMILFGEEDVRVANLKPRISFNFFVTAPKNVSDIIPRTEVEKAIRMSRSRINDAFRLNDNSLEFLGIQPTLGPPNQPPVSPPVATVSKGEEDNMAIIKEFMRFKVHMEGSVNGHEFEIEGEGEGRPYEGTQTAKLKVTKGGPLPFAWDILSPQFMYGSKAYVKHPADIPDYLKLSFPEGFKWERVMNFEDGGVVTVTQDSSLQDGEFIYKVKLRGTNFPSDGPVMQKKTMGWEASSERMYPEDGALKGEIKQRLKLKDGGHYDAEVKTTYKAKKPVQLPGAYNVNIKLDITSHNEDYTIVEQYERAEGRHSTGGMDELYKPPVATLASKTIVLSVGEATRTLTEIQSTADRQIFEEKVGPLVGRLRLTASLRQNGAKTAYRVNLKLDQADVVDSGLPKVRYTQVWSHDVTIVANSTEASRKSLYDLTKSLVATSQVEDLVVNLVPLGRADPLASKTIVLSVGEATRTLTEIQSTADRQIFEEKVGPLVGRLRLTASLRQNGAKTAYRVNLKLDQADVVDSGLPKVRYTQVWSHDVTIVANSTEASRKSLYDLTKSLVATSQVEDLVVNLVPLGRHHHHHH | protein | 140.77 |
| PCP-8x | gagaaacguuucgacauuauauggaaugcgaaacacggaggaugcgggaaacaugaagaucacccauguucgcuuaaccauggauagggaucacccauguugcgguggugcgucaaccagagauuucauaugggaaacucugggacacgcuguauuuauacaugaggaucaccaugugugcuuaaauaugggguaauuccaguuuauauggaaacggaauuaggcuagagcauggcgguauugaguucggguuugaaacgacaauauauggauugcguuuuggggcacgccgucuggaaaggagauuauaugaaaucccuuugcgcgcaaccgguagauacgagucaauauggugaccguaccuugguggggaagcgauaagcacauuauaaggaauggcuuagugcuagacccuggucuuuucgagaaaauaugguuuccgaaaauggguggagguagagggauucucgcgagaag | slncRNA | 154.8 |
| PCP-14x/MCP-15x | gagaaacguuucgacauuauauggaaugcgaaaguggaacguaauggacaugaagacgauuacgcuucacacggaggaugcgggaaacaugaagaucacccauguucgcuuaaccauggauagggaucacccauguugcgguggugcgucaaccagagauuucauaugggaaacucugggacacgcuguauuuauacaugaggaucaccaugugugcuuaaauaugggguaaguugaccauuaggcaacuguaagaugcuccgguuaauuccaguuuauauggaaacggaauugauguaccguugagcaagaacacgauuacgguucuucgauuagauaugguuuaaacgacaauauauggauugcguuuuggggcacgccgucuggagaagaccauuaggcuucuuuacugcgaccgcaauaaaaggagauuauaugaaaucccuuugcgcgcaaccgguagaagaucaccauuagggaucuguaacucacggcgcuauuacgagucaauauggugaccguaaagcuagggcaugugccagaagagcauuagccuucuccuugguggggaagcgauaagcacauuauaaggaauggcuuaaaguggugcggcgggggacuugaccauuaggcaagugugcuagacccuggucuuuucgagaaaauaugguuuccgaaagaacuauacgaagugacaugcgaggauuacccgcauauggugcaaaugggagaauuggaguaaauaugguuacccaauaggcuagagcaugacggcagugagaauuauccacugguuagcggguuaccgagauugcacauuauauggaauggcaauugauucaugccggucguuugugaggaguacccacaaaaugggagggugcuauauaaccagguuauaugcaaccgguuaggccguuguguuaguuuaguucagcauuagcgaacugugcaagacccgguggcuaaggaguuuauauggaaacccuuagcccucgagcaugcuaacaugaggauuacccaugugaugggguuugaaacgugcaauuauuugauauggcaaaaauuggguagggcauggcugcagcgugagaauuauccacgcuaggcuagagcauggcgguauugaguucggguuugagaugaccauuaggcaucugugcuagagcaugcgaaaacgacauaauaugguaugcguuuuggggugcuaguauaccacaugaugagcacacauguauggguggagguagagggauucucgcgagaag | slncRNA | 410.18 |

**Table S1: Sequences of molecular components used in the study, Related to Fig. 1-4.**
